# Atomistic basis of opening and conduction in mammalian inward rectifier potassium (Kir2.2) channels

**DOI:** 10.1101/642090

**Authors:** Eva-Maria Zangerl-Plessl, Sun-Joo Lee, Grigory Maksaev, Harald Bernsteiner, Feifei Ren, Peng Yuan, Anna Stary-Weinzinger, Colin G. Nichols

## Abstract

Potassium ion conduction through open potassium channels is essential to control of membrane potentials in all cells. To elucidate the open conformation and hence the mechanism of K^+^ ion conduction in the classical inward rectifier Kir2.2, we introduced a negative charge (G178D) at the crossing point of the inner helix bundle (HBC), the location of ligand-dependent gating. This ‘forced open’ mutation generated channels that were active even in the complete absence of phosphoinositol-4,5-bisphosphate (PIP_2_), an otherwise essential ligand for Kir channel opening. Crystal structures were obtained at a resolution of 3.6 Å without PIP_2_ bound, or 2.8 Å in complex with PIP_2_. The latter revealed a slight widening at the HBC, through backbone movement. Molecular dynamics (MD) simulations showed that subsequent spontaneous wetting of the pore through the HBC gate region allowed K^+^ ion movement across the HBC and conduction through the channel. Further simulations reveal atomistic details of the opening process and highlight the role of pore lining acidic residues in K^+^ conduction through Kir2 channels.

## Introduction

Potassium channels play essential roles in stabilizing membrane potentials and control of numerous physiological phenomena, in all cells. Inward rectifier (Kir) potassium channels are a major sub-family of channels that lack the classical 4-helix voltage sensing domains of voltage-gated channels, but show voltage-dependence of conductance, due to block by intracellular Mg^2+^ and polyamines at positive membrane potentials, and are controlled by gating in response to regulatory ligand binding (Hibino et al, 2010; Nichols & Lopatin, 1997). Each Kir subfamily exhibits distinct ligand-gating; Kir1 and Kir4/5 are controlled by pH, Kir3 by Gβγ proteins, Kir6 by ADP/ATP and sulphonylurea receptor subunits (SURs)(Nichols & Lopatin, 1997). Phosphoinositol-4,5-bisphosphate (PIP_2_), binding to a canonical PIP_2_ binding site (‘primary’ site) is essential for activation of all Kir channels(D’Avanzo et al, 2010; Hansen et al, 2011; Hilgemann & Ball, 1996). Bulk anionic lipids (PL(-)s) are additional positive allosteric regulators that substantially increase PIP_2_ sensitivity of the Kir2 subfamily(Cheng et al, 2011; Lee et al, 2013). While several Kir crystal(Hansen et al, 2011; Lee et al, 2016; Tao et al, 2009; Whorton & MacKinnon, 2011; Whorton & MacKinnon, 2013) and recently single particle cryo-EM(Martin et al, 2017) structures have been determined, they do not encompass the full conformational ensemble of functional states of the channel. In particular, the M2 (S6) helix bundle crossing (HBC), located close to the cytoplasm-inner leaflet interface, forms a constriction in all available structures that is too narrow for permeation and therefore indicates a closed state of the channel. Lack of open-state structures significantly limits our understanding of the molecular mechanisms by which PIP_2_ and other ligands actually open the channel, of how pore blockers or other ligands actually influence ion conductance, and the details of conductance itself.

A general barrier to progress in molecular level understanding is presented by the potentially obscure functional state of structural conformations determined by X-ray, single particle cryo-EM, or NMR approaches. Molecular dynamics (MD) simulations can provide key functional interpretation of experimentally determined structures. For example, recent MD simulations have revealed the importance of ‘de-wetting’ as the basis of closure at narrow hydrophobic restrictions, and the likelihood that specific structures indeed represent open, conductive states(Aryal et al, 2015; Trick et al, 2015), but close feedback between simulation and experiment is frequently lacking.

We recently obtained crystal structures(Lee et al, 2016) of chicken Kir2.2[K62W] mutant protein (KW) in the absence of, or in complex with, PIP_2_. The K62W mutation endows the equivalent allosteric effect as PL(-) binding, generating physiological sensitivity to activatory PIP_2_(Cheng et al, 2011). The Apo-K62W structure (in the absence of PIP_2_), maintained a high affinity PIP_2_ binding site, revealing a ‘pre-open’ state stabilized by PL(-) binding. However, even after satisfying both of the recognized lipid requirements for opening, the PIP_2_-bound K62W structure was still sterically closed at the inner helix bundle crossing (HBC). This suggests that the open state stability is relatively low and perhaps unlikely to be captured in such crystals.

As an alternate approach to capture the ‘open’ conformation of eukaryotic Kir channels, in the present study we have now introduced an additional mutation (G178D) near the HBC gate that functionally stabilizes open channels, following an approach previously used to obtain an apparently open KirBac3.1 crystal structure(Bavro et al, 2012) and to generate gain-of-function mutants of various other K channels(Brelidze et al, 2003; Nimigean et al, 2003), and have determined crystal structures of these mutant channels with or without bound PIP_2_. The PIP_2_ bound KW/GD structure was slightly more open (by ~1.5 Å diameter) at the HBC than the previous PIP_2_-bound K62W structure(Lee et al, 2016). Molecular dynamics simulations reveal that, when embedded in a lipid membrane, relaxation of this crystal structure, together with side-chain flexibility, results in wetting of the pore through the HBC region. This leads to rapid spontaneous further opening of this region and K^+^ conductance through the entire channel, revealing novel details of the ion permeation process.

## Results

### Generation of a ‘locked open’ Kir2.2 channel

In order to force Kir2.2 channels into an ‘open’ conformation, we have followed the approach first used to obtain an apparently open KirBac3.1 structure(Bavro et al, 2012), introducing a charge mutation near the helix bundle crossing (HBC) gate in each subunit. By electrostatic repulsion, such introduced charges may provide an energetic ‘push’ against one another to stabilize the open state. We introduced G178D (Supplementary Fig. 1a,b) on both the truncated wild-type chicken Kir2.2(Tao et al, 2009) as well as the Kir2.2[K62W] mutant background(Lee et al, 2016). Purified channel proteins were incorporated into liposomes and characterized functionally by ^86^Rb^+^ flux assays. As shown previously(Lee et al, 2016), both WT Kir2.2 and Kir2.2[K62W] (KW) channels were inactive in 9:1 POPE:POPG, and required 0.1% PIP_2_ for activity. In contrast, both single mutant Kir2.2[G178D] (GD) and double mutant Kir2.2[K62W, G178D] (KW/GD) channels were active in the complete absence of PIP_2_ (Supplementary Fig. 1c), although KW/GD showed further stimulation of activity when PIP_2_ was present. For unknown reasons, the protein yield of the single GD mutant protein was consistently very low, and we therefore focused further analysis on the KW/GD double mutant proteins.

**Fig 1.**
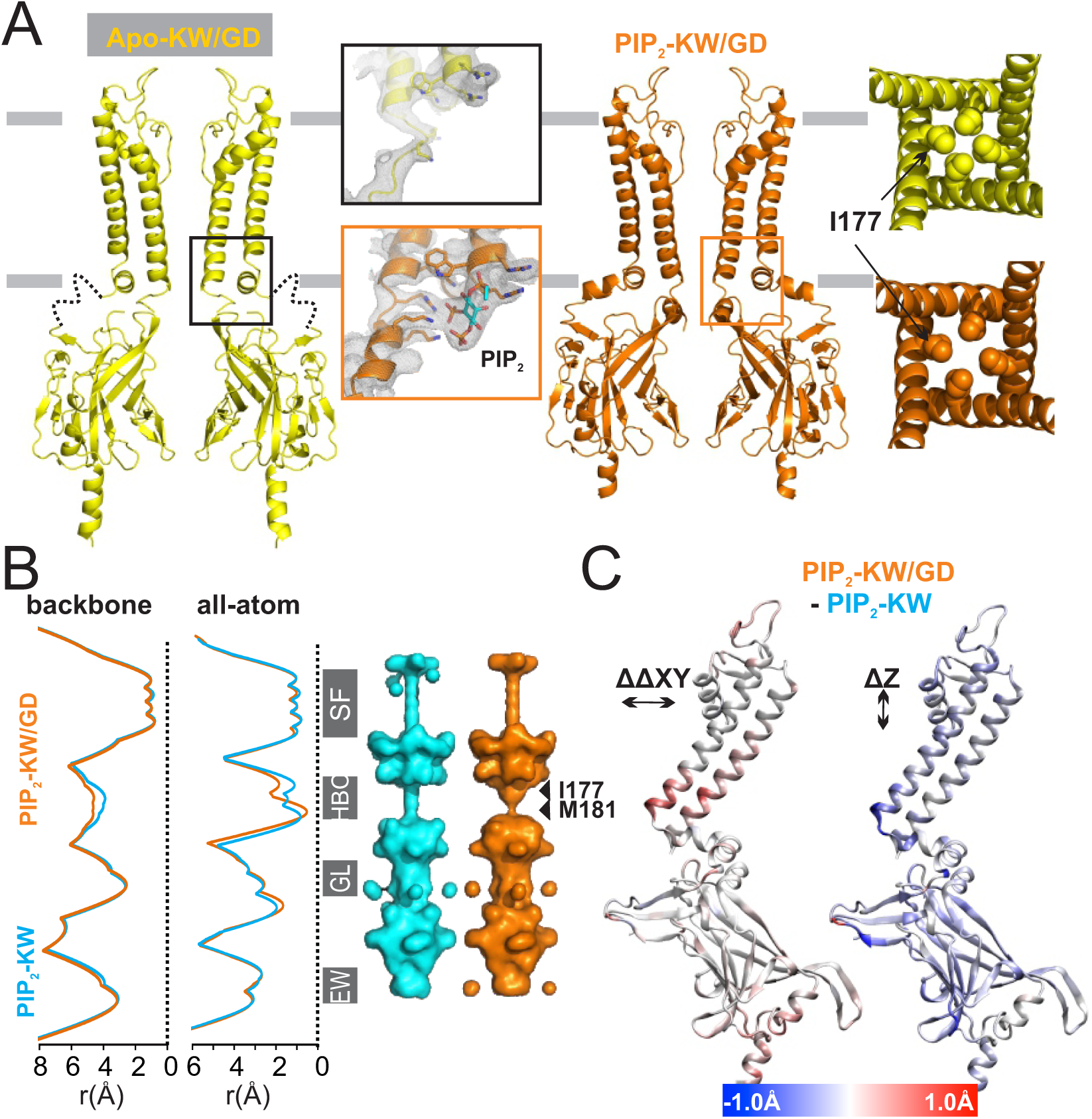
Crystal structures of forced open mutant channels. **A**) (*left*) Apo-KW/GD (yellow) and PIP_2_-KW/GD (orange) crystal structures are shown in ribbon diagram. For clarity, only two subunits are shown for the side view. The local conformations of the PIP_2_ binding site region are shown in the inset, with 2fo-fc electron density contoured at 1 σ. (*right*) A ‘top down’ view of each structure in the plane of the membrane shows the HBC region with I177 side chains depicted in balls. **B**) Pore-radius profiles of PIP_2_-KW (cyan) and PIP_2_-KW/GD (orange) crystal structures computed by HOLE2 are shown with backbone atoms (left) or with all atoms including side chain (right). Pore-lining surfaces of the PIP_2_-KW (cyan) and PIP_2_-KW/GD (orange) crystal structures are shown with the major constrictions designated: SF: selectivity filter, HBC: helix bundle crossing (I177, M181), GL: G-loop, and EW:entry way (vertically aligned to the ribbon diagrams in C). **C**) Diagonal Cα distance changes (ΔΔCα = PIP_2_-KW/GD(ΔCα) – PIP_2_-KW(ΔCα)) in the lateral dimension (the xy plane) (left) and vertical movement along the z axis of Cα atoms (right) mapped on a single Kir2 subunit. The color bar shows the ranges of distance change.

Wild type Kir2 channels are specifically activated by PIP_2_, while other naturally occurring phosphoinositides tend to show inhibitory effects(D’Avanzo et al, 2010). Sensitivity to PIP_2_ was similar for KW and KW/GD channels, with apparent Kd of ~0.03 (w/w)%, which corresponds to ~0.02 mol% in each case. Neither channel was sensitive to PI(4)P_1_, at up to 1 (w/w) %, corresponding to ~0.7 mol% -, indicating that phosphoinositide specificity was not altered by the ‘forced open’ G178D mutation (Supplementary Fig. 1d). In these ^86^Rb flux assays, ion selectivity and spermine(Lopatin et al, 1994) block were also indistinguishable between WT, KW, and KW/GD, with or without PIP_2_ (Supplementary Fig. 1e,f), indicating that, whether separately or in combination, the G178D mutation does not markedly alter sensitivity to regulatory ligands, other than increasing the relative stability of the channel open state.

### Crystal structures of Apo-KW/GD and PIP_2_-KW/GD

KW/GD mutant crystals were obtained in both the absence (Apo-KW/GD) and presence (PIP_2_-KW/GD) of di-oleoyl phosphoinositol-(4,5)-bisphosphate (18:1 PIP_2_). Crystal growth conditions were similar, although high concentrations (>130 mM) of triNaCitrate were required for the growth of Apo crystals, while lower triNaCitrate concentrations (<100 mM) facilitated PIP_2_-KG/WD crystal formation. X-ray diffraction data were collected at 3.6 Å resolution, and a final structural model was obtained with R value of 23.7 % and R_free_ value of 29.1 % for Apo-KW/GD. PIP_2_-KW/GD crystals diffracted to 2.8 Å, and the structural model was refined with R value of 22% and R_free_ value of 27% (Fig. 1, see Methods, and details of crystals and models in Supplementary Table 1). In both proteins, the G178D side chains face the inner helix of the neighboring subunit (Supplementary Fig. 2), rather than pointing directly into the pore axis. However, an obvious difference between Apo-KW/GD and PIP_2_-KW/GD structures is that the CTD is disengaged from the TMD in the Apo-KW/GD structure (‘loose’ conformation), with an unstructured and unoccupied PIP_2_ binding site, similar to the Apo-WT structure (PDB ID:3JYC)(Tao et al, 2009), whereas the CTD is tightly engaged with the TMD in the PIP_2_-KW/GD structure (‘compact’ conformation), with strong PIP_2_ density in the fully formed binding site (Fig. 1A, inset).

As in previous Kir2.2 structures, the disengaged Apo-KW/GD structure is closed and non-conducting, due to a steric constriction throughout the HBC region (Fig. 1A, right). The PIP_2_-KW/GD structure (Fig. 1A, right) is subtly, but noticeably, different from all other structures, including the counterpart PIP_2_-bound KW crystal structure (PIP_2_-KW, 5KUM(Lee et al, 2016)) at the HBC location. Superimposed PIP_2_-KW and PIP_2_-KW/GD crystal structures show that structural changes due to the GD mutation are localized near the HBC, resulting in a minor (~ 1 Å) lateral expansion at the bottom of the TM1 and TM2 helices (Fig. 1B,C). Pore dimensions analyzed by HOLE2 (Smart et al, 1996) and HOLLOW(Ho & Gruswitz, 2008) show a widening at residue 177 due to this M2 helix backbone displacement (Fig. 1B). The pore is still narrowed immediately below this by M181 sidechain projections (Fig. 1B), but flexibility in the M181 sidechain could lead to rapid widening of this region. Since hydration or wetting of hydrophobic residues that form the gate is a prerequisite for ion channel opening(Aryal et al, 2015; Trick et al, 2015), the slight expansion at the backbone might then be sufficient to allow water permeation at the HBC (Fig. 1B,C) and thereby render the channel ‘open’. To examine this possibility, we carried out all-atom molecular dynamic (MD) simulations of the PIP_2_-KW/GD crystal structure (see Methods). Unlike MD simulations of previous Kir2.2 crystal structures(Lee et al, 2016), which stayed closed unless external force was applied to pull open the HBC gate(Li et al, 2015), the TM helices and M181 sidechains spontaneously moved outwards, resulting in HBC gate expansion of ~5 Å (Fig. 2A) and HBC ‘wetting’ in all PIP_2_-KW/GD simulations. This in turn led to permeation of potassium ions through the HBC gate as well as the SF, at rates in a similar order to those measured experimentally, as described in detail below.

**Fig 2.**
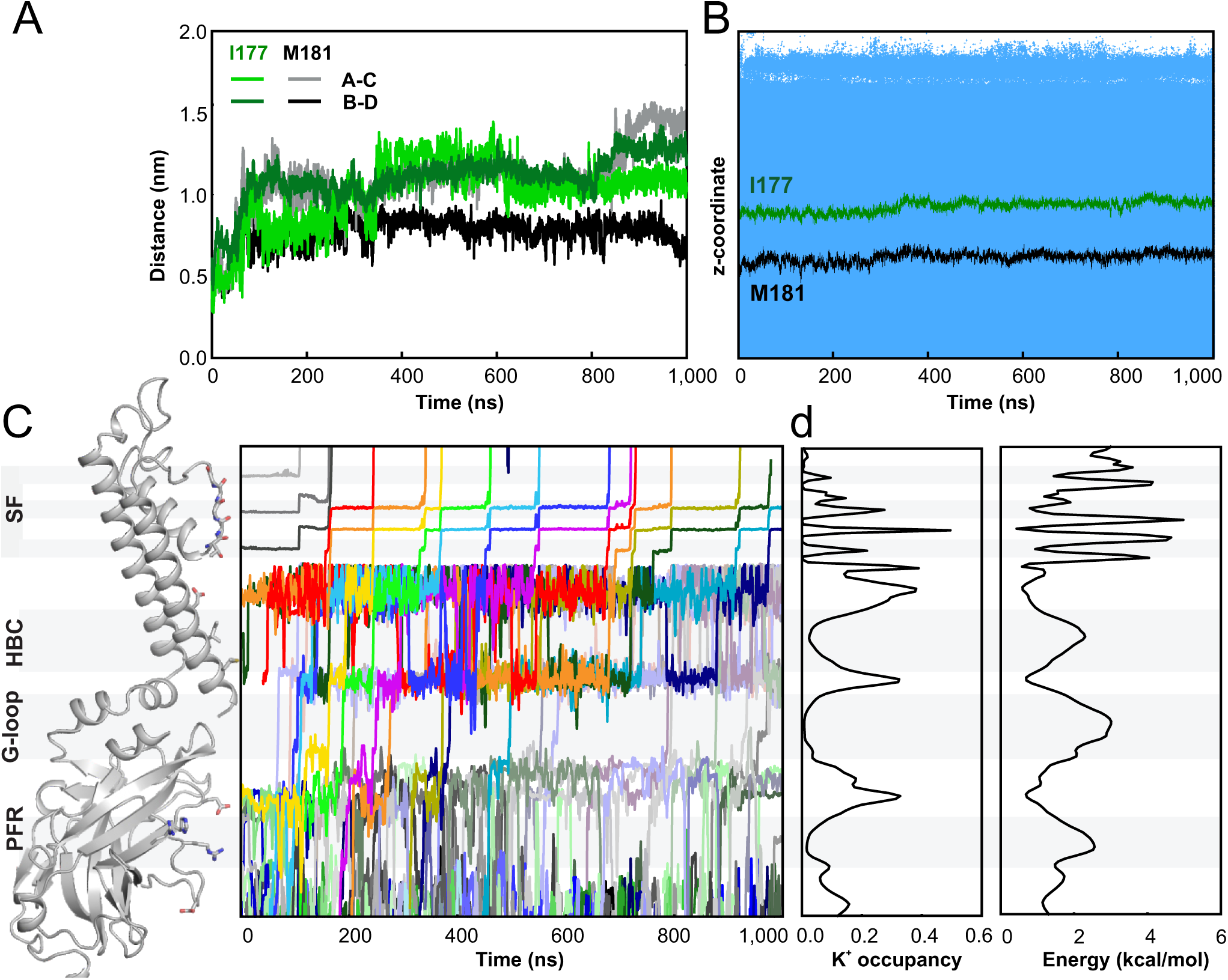
Ion flux through open conducting KW/GD structures. **A**) The evolution of minimal distances of residues I177 and M181 between two opposing subunits over the course of 1 µs simulation. **B**) Location of water molecules (represented as small blue dots) within the pore near the HBC gate over the course of 1 µs simulation. The HBC gate location is indicated by the I177 (green) and M181 (black) Cα positions. **C**) Time trajectories of individual K^+^ ion (varying colors) are shown from the entry way of the pore to beyond the S0 position in the SF (vertical positions match the ribbon diagram on the left). **d**) (*left*) Ion occupancy along the pore averaged over all simulations (total simulation time of 3.8 μs), from which the free energy profile for potassium ions along the pore axis is obtained (*right*) via an inverse Boltzmann transformation of the K^+^ ion density profile.

### K^+^ ion conduction through the open channel

To gain insight to the permeation mechanism through open Kir2 channels, we performed repeated simulations (for details see Supplementary Table 2) of the PIP_2_-KW/GD structure. Channel opening resulted from spontaneous expansion at the HBC (Fig. 2A) and consequently uninterrupted water accessibility through the entire HBC region (Fig. 2B). In all (6 out of 6) simulations, the HBC opened within 50 ns and, following the first 50 ns of simulation, the shortest diagonal distances at the HBC measured on average 6.3 ± 1.3 Å at the I177 site, and 10.7 ± 2.9 Å at M181. Fig. 2C illustrates the progressive location of individual ions through the pore during one long (1,000 ns) simulation. During such simulations, K^+^ ion distributions within the channel reached essentially steady– state, allowing assessment of the mean location of K^+^ ions in the open, conducting channel. During permeation, K^+^ ions were highly localized near the negative charges that underlie weak rectification (E225) (on average 3.4, maximum 5) within the CTD, as well as near the ‘rectification controller’ negative charges (D173) (on average 4.7, maximum 6) in the inner cavity. They were also localized near the introduced D178 (on average 3.2, maximum 5) at the HBC gate (Fig. 2C). K^+^ ion distributions (Fig. 2d left), averaged over the entire 2.8 μS of simulation, reveal effective energy barriers at the G-loop, the HBC and the SF (Fig. 2D). These barriers are ~3 kcal/mol and 1.9 kcal/mol at the G-loop and the HBC respectively (Fig. 2D, right), but both are relatively low and suggest that these locations will not function as significant rate-limiting steps for K^+^ ion permeation. More substantial energy barriers in the SF (4.3 kcal/mol) are likely to provide major rate-limiting steps for K^+^ conduction although, in absolute terms, these are still only slightly higher than what would be expected for diffusion-limited ion flux(Berneche & Roux, 2001), and hence will not preclude high K^+^ conductance through the channel.

These simulations illustrate elementary steps underlying outward movement of K^+^ ions through the channel (Fig. 2C). Ions enter the cytoplasmic pore from the intracellular side by free diffusion, with no preferred path, until they reach a narrow region, which we will term the Entry Way (EW), formed by the short helices (F^255^DKG^258^, see Supplementary Fig. 3), which obliges the ions to pass in single file to reach the ring of negative charges at E225. In between these two regions, positively charged amino acids (H227, R229 and R261) form a “positive focusing ring” (PFR) (Fig. 2C, see Supplementary Fig. 3) that prohibits more than one positively charged ion at a time from passing through. Multiple ions are coordinated by the E225 residues, but ions then transition in single file to the region between the G-loop gate and the HBC gate, where up to 5 ions are again coordinated by the G178D residues. The ions pass the HBC in a fully solvated state (see Supplementary Fig. 4), but in single file, to reach the inner cavity (for a full conduction event see Supplementary Movie 1). As discussed above, major energy barriers are present in the selectivity filter, in particular between the S3 and S2 positions (Fig. 2D), leading to residence times for K^+^ ions in S3 of 8-131 ns. Nevertheless, in the two longest (1,000 ns) simulations, 14 and 6 ions, respectively, passed completely through the channel, while in the shorter (200ns) simulations, 6, 4, 1 and 0 ions passed through. With an applied transmembrane field of 584mV, this corresponds to a conductance of up to 8 pS. Although lower, this is a similar order of magnitude to experimentally measured open Kir2.2 channel conductance (~35 pS in symmetric 140 mM [K^+^](Takahashi et al, 1994)).

### K^+^ conduction through open ‘wild-type’ KW/GD(G) channels

To examine whether the conducting state results from the steric widening that occurs in the PIP_2_-KW/GD crystal structure, or is a direct electrostatic effect of the introduced G178D negative charges, we carried out additional simulations on channels that were back-mutated *in silico* at residue 178 from Asp to the native Gly (KW/GD(G)), after 50 ns pre-simulations of the PIP_2_-KW/GD crystal structure, i.e. after initial wetting of the HBC gate had occurred. Importantly, wetting was subsequently maintained, and channels continued to conduct, over the full length of the subsequent simulations. In total, 10 replica simulations of 200 ns and one of 1,000 ns length were carried out. In all cases, although the minimum distances at the HBC were slightly narrower than in PIP_2_-KW/GD simulations at the level of residue 177, the PIP_2_-KW/GD(G) structures remained open and conducting in these simulations (Fig. 3; For a full conduction event see Supplementary Movie 2). In these PIP_2_-KW/GD(G) simulations, K^+^ ions still localized near residues E225 (average 3.5, maximum 5) in the CTD and residues D173 (average 3.7, maximum 5)) in the inner cavity, but now ions were much less likely to localize near residue 178 (average ~1, maximum 2). Otherwise, ion distributions in, and ion fluxes through, the KW/GD(G) channel were quite similar to KW/GD (Fig. 3C,D). K^+^ ion distributions averaged over the total 3,000 ns of simulation revealed energy barriers at the G-loop and SF that were similar to those in KW/GD (Fig. 3D, left), but the slightly narrower opening of ~6.3 Å and concomitant occasional de-wetting phases (Fig. 3B), at the HBC resulted in a slightly higher free energy barrier at the HBC gate (3.5 kcal mol^−1^, c.f. 1.9 kcal mol^−1^ in KW/GD, Fig. 3D, right), and slightly lower net conductances (up to 5.5 pS).

**Fig 3.**
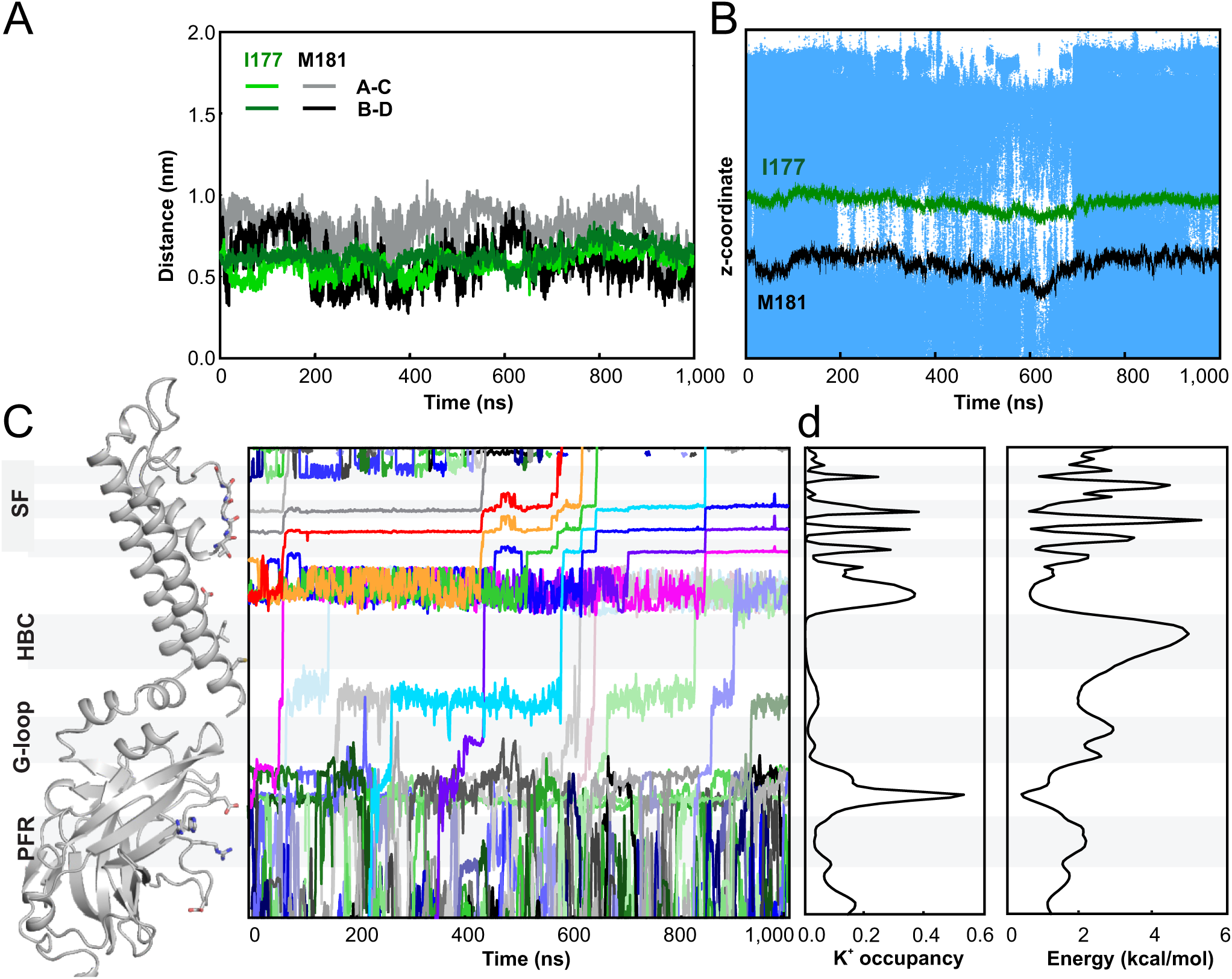
Ion flux through KW/GD(G) open conducting structures. Corresponding information as in Fig. 2 for KW/GD(G) simulations (composed of 10 replica simulations of 200 ns as well as one 1μs simulation). **A**) The evolution of minimal distances of residues I177 and M181 between two opposing subunits over the course of 1 µs simulation. **B**) Location of water molecules (represented as small blue dots) within the pore near the HBC gate over the course of 1 µs simulation. The HBC gate location is indicated by the I177 (green) and M181 (black) Cα positions. **C**) Time trajectories of individual K^+^ ion (varying colors) are shown from the entry way of the pore to beyond the S0 position in the SF (vertical positions match the ribbon diagram on the left). **D**) (*left*) Ion occupancy along the pore averaged over all simulations (total simulation time of 3.0 μs), from which the free energy profile for potassium ions along the pore axis is obtained (*right*) via an inverse Boltzmann transformation of the K^+^ ion density profile.

### The structural basis of conducting states

As a consequence of decreased ion occupancy in the pore, the above simulations indicate lower conduction for KW/GD(G), in which the ‘forced open’ channel is ‘back mutated’ to a wild type pore without the four acidic Asp residues at the HBC gate, than for KW/GD channels. We experimentally assessed the role of the introduced Asp residues in channel conductance, with single channel recordings of KW/GD and KW proteins expressed in Cosm6 cells (Fig. 4A,B). Even though absolute unitary conductance levels predicted above by MD simulations were lower than the experimentally measured values, the relative effect of the introduction of acidic Asp residues at position 178 was strikingly similar: the KW/GD mutant channel conductance was ~30% higher than that of WT channels (Fig. 4A,B), similar to the ~40% higher conductance of KW/GD in the simulations.

**Fig 4.**
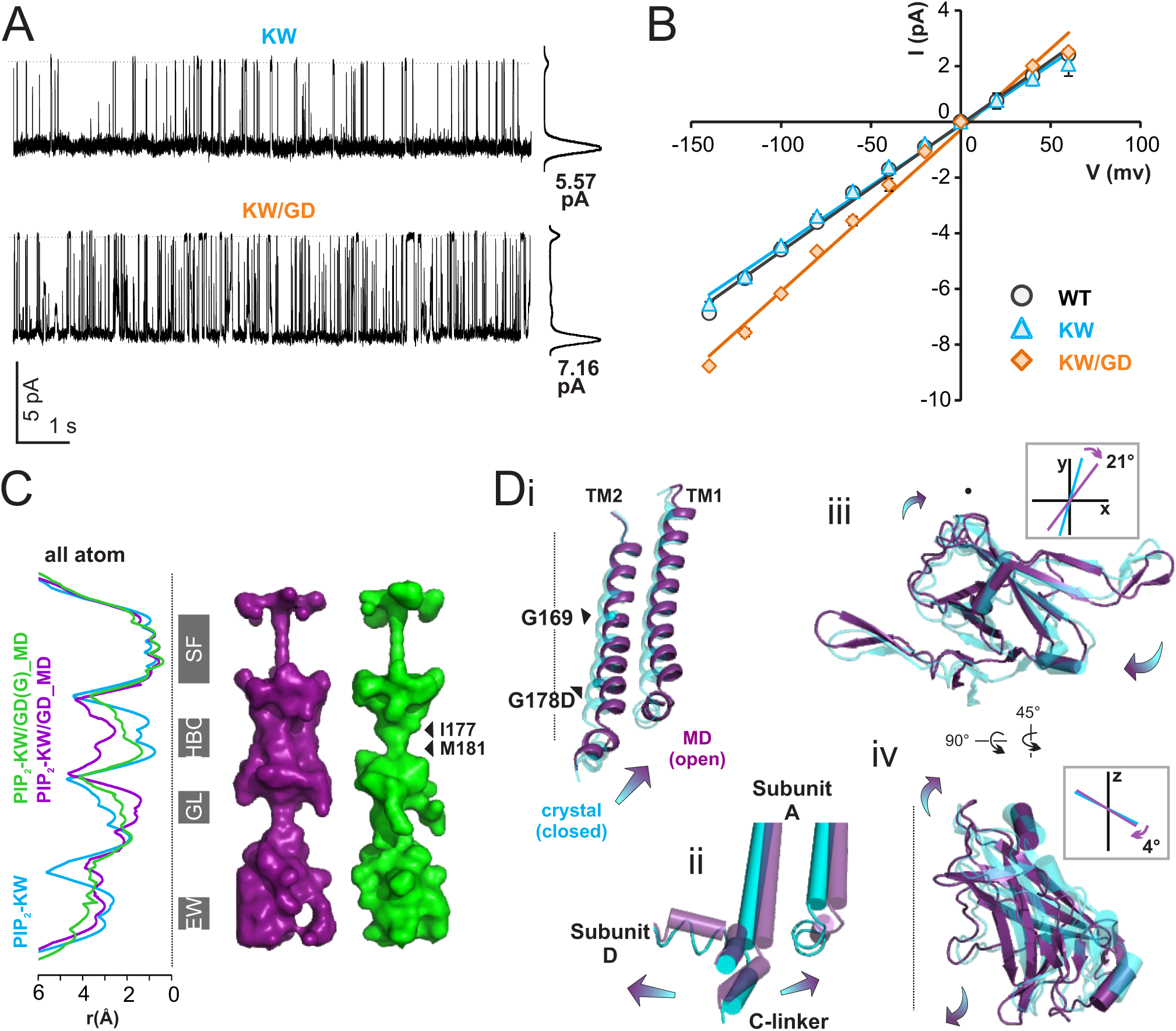
Kir2 channel unitary conductances and open structures. **A**) Representative single-channel currents recorded from CosM6 cells at −120 mV and 150 mM symmetrical K^+^ condition using inside-out patch configuration. **B**) Single-channel current-voltage relationships. Symbols and error bars are mean and standard deviation (n => 4), and the lines are linear fit of the data. **C**) The last snapshots of 1 µs MD simulations of PIP_2_-KW/GD and PIP_2_-KW/GD(G) were compared to the closed PIP_2_-KW crystal structure (5KUM) in order to assess the open state structures. (*left*) Pore-radius profiles of PIP_2_-KW/GD_MD (purple), PIP_2_-KW/GD(G)_MD (green), and PIP_2_-KW (cyan) were computed by HOLE2 with all atoms including side chain. (*right*) The pore-lining surfaces of PIP_2_-KW/GD_MD (purple) and PIP_2_-KW/GD(G)_MD (green) computed by HOLLOW are shown with the major constriction sites designated: SF: selectivity filter, HBC: helix bundle crossing (I177, M181), GL: G-loop, and EW: entry way. **D**) Closed PIP_2_-KW crystal structure (cyan) and open PIP_2_-KW/GD_MD (purple) structure are overlaid; arrows indicate predominant conformational changes from closed to open state. (i): TM1 and TM2 helices are shown in ribbon diagram. (ii): cylindrical cartoon representation of subunit interface. (iii): a top-down view of CTD domains in ribbon diagram. The central vectors of the C-linker helices projected on the xy-plane are overlaid to show a twisting motion in the inset. (iv): side view of CTD domain in plane of the membrane normal. The central vectors of the C-linker helices projected on the z-axis are overlaid to show a tilting motion in the inset.

As representative open state structures, the last snapshots of 1,000 ns PIP_2_-KW/GD and PIP_2_-KW/GD(G) MD simulations are compared to the PIP_2_-KW crystal structure in Fig. 4C,D. The conformational changes that generate a conducting channel are qualitatively similar in both KW/GD and KW/GD(G): the prohibitive narrowing at residue 181 is removed in both, but the HBC region is considerably wider in the former (Fig. 4C). TM2 helix bending brings about slight upward and counterclockwise conformational translation (Fig. 4D i): bending at the highly conserved glycine hinge (residue G169) with average bending values of 14.7° ± 10.1° for PIP_2_-KW/GD and 10.9° ± 8.0° for PIP_2_-KW/GD(G) and additional helix bending at G178 location, with average values of 9.5°± 5.4° for PIP_2_-KW/GD and 6.5° ± 4.3° for PIP_2_-KW/GD(G), computed by VMD Plugin Bendix(Dahl et al, 2012).

A top-down view of these snapshots (Fig. 4D iii) reveals CTD motions in conducting channels relative to the crystal structure (highlighted by morph video based on PIP_2_-KW crystal structure and KW/(GD)G MD open structure, Supplementary Movie 3). These include clockwise whole CTD rotational motions about the central pore axis of 4.7° ± 1.6° for KW/GD and 1.45° ± 2.25° for KW/GD(G) simulations. More noticeable was twisting of individual subunit CTDs around an axis within each subunit, with the pivot point at the c-linker (Fig. 4D iii). A side view of the CTD reveals additional tilting motions that pull the top of the CTD away from the pore axis, while the bottom of the CTD is pulled toward the pore axis (Fig. 4D iv) in the conducting channel.

### Direct knock-on permeation

The inner cavity contained 4-5 ions, localized at the level of residues D173 (the ‘rectification controller’) and T143, towards the top of the selectivity filter (see Figs. 2C, 3C) during steady-state KW/GD or KW/GD(G) simulations. Following initiation of each KW/GD simulation, with one K^+^ ion placed in the inner cavity, transitions from the inner cavity to selectivity filter site S4 (formed by the carbonyl oxygen atoms of T143), did not begin until after the four D173 charges were balanced by four K^+^ ions. Once an ion entered the SF site S4, the ion in site S3 was rapidly pushed via a direct knock-on upwards to S2, leading to a rapid exit of the ion initially located at site S2 to the extracellular side, via S1 and S0 (Fig. 5, see Supplementary Movie 4). Thus, while initial conditions were set up with K^+^ ions in sites S0, S2, and S4, and water molecules in sites S1 and S3, S2 and S3 were always rapidly and preferably occupied by K^+^ ions (see Fig. 5), leaving sites S0, S1, and S4 filled with water molecules. Importantly, except for the water molecules initially placed in S1 and S3, no additional water entered or permeated the selectivity filter throughout the simulations. Thus, our simulations implicate direct knock-on as the underlying mechanism for high conductivity of K^+^ ions through the channel (Fig. 5 and Supplementary Movie 4), as also seen in other recent simulation studies (Kopec et al, 2018; Köpfer et al, 2014) (see Discussion).

**Fig 5.**
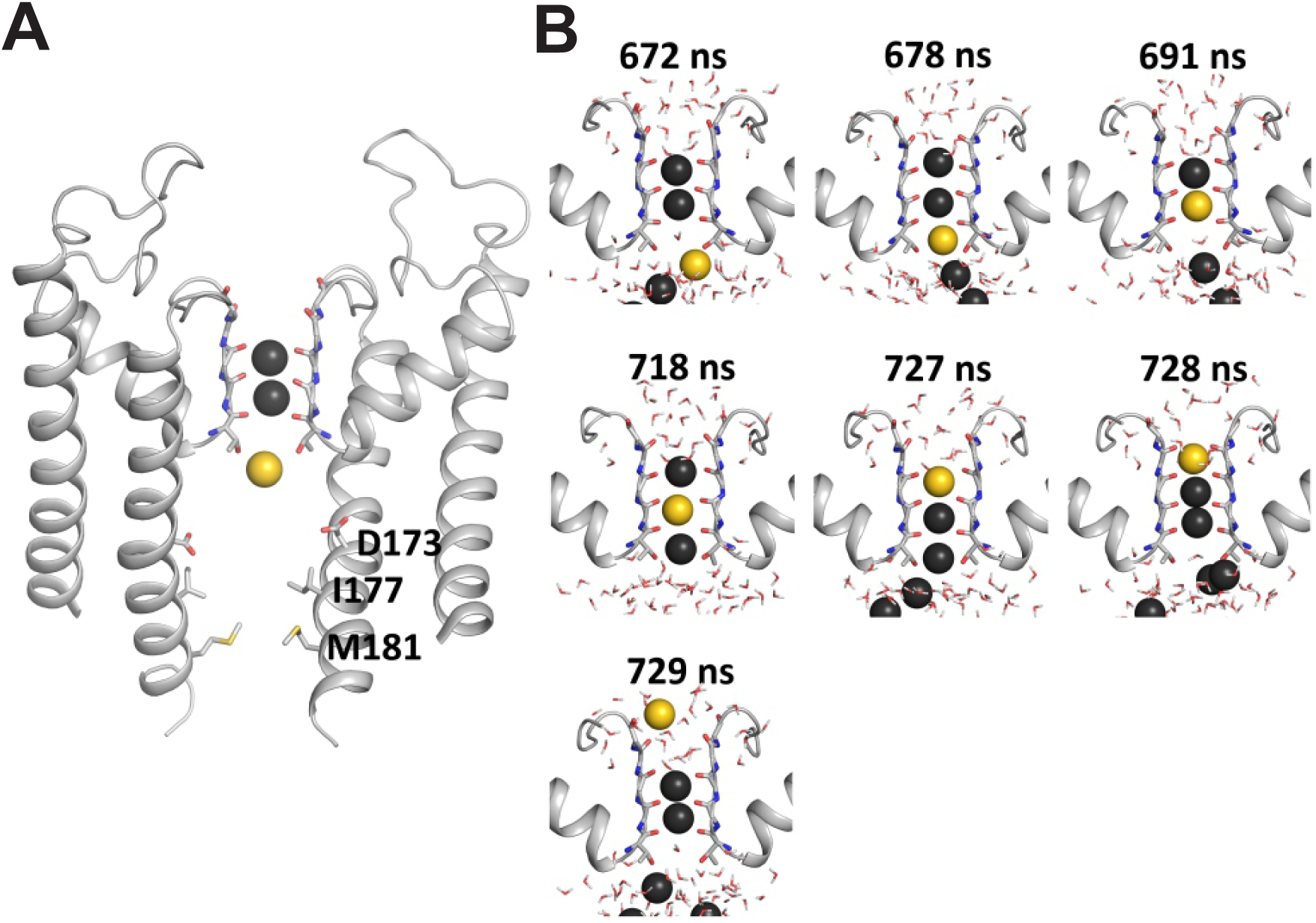
Direct knock-on K ion movement in selectivity filter. **A**) Pseudo-steady state conformation of the SF with two resident K^+^ ions (black) in S2 and S3, immediately before permeation event initiated by entry of inner cavity K^+^ ion (yellow). For the selectivity filter, the backbone of residues IGYG (residue numbers 144 to 147) as well as the full T143 are shown as sticks as well as the rectification controller residue D173. For clarity, only two opposing domains are shown. **B**) Snapshots of the selectivity filter during the following conduction event. Surrounding water represented as sticks. Consecutive simulation snapshots show the complete permeation of the yellow K^+^ ion, followed by additional K^+^ ions (black) from the inner cavity, with no intervening water molecules.

## Discussion

### The mechanism of Kir channel opening

K channels gate in response to channel-specific stimuli, including water-soluble or membrane-soluble ligands and/or membrane potentials(Grandi et al, 2017). Rather than interacting directly with the pore domain, these stimuli typically interact with or transform regulatory domains, which then effect opening or closing of the pore. In Kir channels, gating is controlled by diverse ligands and regulators (Na^+^, G_βγ_, ethanol, ATP, SURs) in addition to the conserved primary agonist, PIP_2_(Hibino et al, 2010). In each case, these ligands bind to the cytoplasmic domain (CTD) to control opening and closure of the channel through physical coupling to the gate, a steric restriction located at the bottom of the transmembrane pore domain. Multiple high resolution structures of different eukaryotic Kir channel family members have been obtained, but always in a closed or non-conducting state(Hansen et al, 2011; Lee et al, 2016; Tao et al, 2009; Whorton & MacKinnon, 2011; Whorton & MacKinnon, 2013). In an effort to overcome this limitation, we here generated a mutant Kir2.2 channel (KW/GD) that is essentially locked open functionally, and obtained high resolution crystal structures. These structures show a minor expansion at the HBC gate compared to previous Kir2.2 crystal structures, but when placed in a membrane and freed from crystallographic contacts *in silico*, the structures undergo rapid spontaneous expansion at the HBC, enabling water and K^+^ ion flux through the HBC gate as well as through the selectivity filter.

Ligand-dependent movements of the Kir CTD with respect to the TMD have been reported in crystallographic, FRET and simulation studies(Bavro et al, 2012; Whorton & MacKinnon, 2011; Whorton & MacKinnon, 2013). Consistent with these studies(Wang et al, 2012; Wang et al, 2016), Kir channel crystal structures determined in complex with different combinations of ligands suggest that two major conformational changes occur concurrent with channel opening - CTD rotation/twisting and CTD tethering to the TMD. In our MD simulations, we also observe gating-dependent clockwise rotational motions of the CTD relative to the TMD in comparison to crystal structures. However, these CTD rotational motions are both relatively small and bidirectional. Recent Cryo-EM structures of Na^+^-activated K^+^ channels also show independence of rotational motion of the CTD from the channel gating transition(Hite & MacKinnon, 2017), and it is unclear whether rotation in any particular direction is directly linked to channel opening.

More interestingly, we observed rigid body twisting and tilting of individual subunit CTDs around their own axes with the pivot at the C-linker of each subunit (Fig. 4D). These motions likely result from the tight coupling of the CTD to the TM helices and slide helix, in turn mediated by the C-linker, which functions as a hub between the TM helices and each CTD (Fig. 4D, ii). Counter-clockwise outward and upward bending of TM helices within the same subunit pulls the C-linker in a counter-clockwise direction, while the same TM helix motions in the neighboring subunit to the left pulls the C-linker in a clockwise direction, resulting in twisting and tilting of the C-linker to cause a counterclockwise rotation and up- and outward motion of the top of the CTD while the lower part rotates in the clockwise direction and toward the pore axis. Such rigid body motions are in accordance with changes in FRET between CTDs during KirBac1.1 opening and closing(Wang et al, 2012; Wang et al, 2016), which require subunit CTD motions relative to one another and would not be registered by whole CTD rotational motions about the pore axis, since the latter involve no diagonal distance changes.

Interestingly, the observed intra-subunit CTD motions in our simulations of PIP_2_-dependent Kir2.2 opening are essentially the same as the conformational changes predicted in KirBac1.1 following PIP_2_ binding(Wang et al, 2012), even though these result in channel closure in the latter case. This is consistent with the idea that PIP_2_ binding facilitates the same secondary structural changes in both channels, but that these are coupled (through as yet unclear final pathways) to opposite functional consequences in prokaryotic and eukaryotic Kir channels(Cheng et al, 2011; D’Avanzo et al, 2010). One obvious gating-associated conformational change is tethering of the CTD to the TMD to form a ‘compact’ structure (Hansen et al, 2011; Lee et al, 2016; Tao et al, 2009). The first PIP_2_-bound Kir2.2 structure (PDB ID: 3SPI) suggested that PIP_2_ may be responsible for causing this vertical motion of the CTD by inducing formation of the PIP_2_ binding site from the unstructured Apo-structure [PDB ID: 3JYC]. However, subsequent GIRK2 crystal structures, which adopted a compact conformation even in the absence of PIP_2_ [PDB ID:3SYO](Whorton & MacKinnon, 2011), suggest that other factors could lead to generation of the compact CTD conformation. Our recent crystal structures show that Kir2.2 also adopts the compact conformation in the absence of PIP_2_, if the requirement for additional tethering of the CTD to the membrane at a second, low affinity, PL(-) site(Cheng et al, 2011; Lee et al, 2013) is met(Lee et al, 2016). Thus the primary effect of PIP_2_ binding, rather than being to induce the compact conformation, may be to stabilize the compact conformation, thereby facilitating additional opening movements.

### The number of ions in the pore

While all K channels exhibit specificity for K^+^ ions over Na^+^ ions, each K channel subfamily is unique in single channel properties(Naranjo et al, 2016), with conductance of the single channel being dependent on rate-limiting structures within the permeation path. Multiple studies of various K channel sub-types have converged to the same conclusion that negative charges lining the pore below the SF as well as steric dimensions (width and length) of the hydrophobic regions of the pore act as major determinants of single channel conductance, by varying K^+^ ion accessibility to the SF(Brelidze et al, 2003; Diaz-Franulic et al, 2015; Naranjo et al, 2016; Nimigean et al, 2003). The Kir channel pore is uniquely long (~7 nm), due to the cytoplasmic extension (~ 4 nm) that is provided by the intracellular CTD(Naranjo et al, 2016), and also lined by several charged residues (3 acidic and 3 basic residues per subunit). Point mutations that reduce the negative charges in the CTD decrease K^+^ conductance, while reduction in positive charges in the CTD increase K^+^ conductance(Fujiwara & Kubo, 2006), implying that local K^+^ concentrations below the SF are critical. Experimentally, we show here (Fig. 4A,B) that introduction of additional negative charges at the residue 178 positions results in higher single channel conductance for the KW/GD mutant, compared to the KW and WT counterparts (Fig. 4A,B). In addition to showing how the D178 negative charges cause conformational changes that generate the ‘open’ conducting state (see above), MD simulations of KW/GD versus the back-mutated KW/GD(G) indicate that the D178 negative charges lower the barrier for conductance through the HBC gate region (1.9 vs. 3.5 kcal mol^−1^) by increasing K^+^ ion concentrations in the inner cavity (compare Figs. 2 and 3) which then increases unitary conductance by increasing the ‘knock on’ pressure for ions to move into and through the selectivity filter.

It is important to note that the above interpretation, i.e. that multiple ions can be simultaneously present at a given depth within the pore, is at least superficially inconsistent with interpretations of crystallographic studies. Multiple high resolution crystal structures indicate that K^+^ ions are vertically aligned along the central pore axis, with essentially no implications that more than one ion can be present simultaneously at any location(Lee et al, 2016; Xu et al, 2009), including within the inner cavity(Tao et al, 2009). A crystal structure of Kir2.2[K62W] at 2.0 Å resolution(PDB ID:5KUM)(Lee et al, 2016) indicates at most two K^+^ ions in the inner cavity (see Supplementary Fig 5). By contrast, the KW/GD MD simulations predict K^+^ ion occupancy in the open conducting Kir2.2 inner cavity as high as 4.7, with 3.5 K^+^ ions continuously present near E225 in the cytoplasmic cavity. It is conceivable that acidic residues within the channel pore may be protonated due to close apposition(Bradley et al, 2012), and predicted pKa values by PDB2PQR/APBS webserver(Gosink et al, 2014) are indeed close to pH 7.0 for D173 and pH 6.0 for E225 in the closed KW and KW/GD structures (Supplementary Table 3). These values are much higher than the pKa values of these residues in solution (~pH 2.0). Thus, at least in the closed channels, not all acidic residues may be charged. However, countering this notion that the formal negative charge of the Kir pore may be very low in closed channel structures, cKir2.2 structures can clearly hold at least 2-3 positive charges in the inner cavity and cytoplasmic cavity based on heavy atom (Sr^2+^, Eu^3+^) anomalous diffraction(Tao et al, 2009), although this might be explained by a decrease in apparent pKa values in the presence of strong positive charges, i.e. the protonation state of the acidic residues may be sensitive to the presence of ion charges and the inner cavity could accommodate different ion charges without undergoing any conformational changes.

Additional experimental findings support assignation of full negative charge to these acidic residues in the MD simulations. First, the Kir6.2 channel inner cavity with 4 introduced acidic charges at the ‘rectification controller’ (N160D) can enclose spermine^4+^ (Phillips & Nichols, 2003), suggesting that four positive charges can cohabitate with and be balanced by four negatively charged aspartic residues. Second, the voltage-dependence of spermine block of Kir6.2 is directly dependent on the number of negative charges in the inner cavity, whether changed by the number of mutant N160D residues, or by adding positive charge via MTSEA or MTSET modification(Kurata et al, 2010). The only straightforward interpretation of such results is that four negative charges in the inner cavity are required for steep spermine-induced rectification and that at least in the open channel, the rectification controller residues are indeed charged.

### The mechanism of K^+^ ion conduction

Mechanisms of ion translocation through the selectivity filter of various voltage gated K^+^ channels, as well as through KcsA, have been interrogated previously using MD simulations(Åqvist & Luzhkov, 2000; Berneche & Roux, 2001; Furini & Domene, 2009; Köpfer et al, 2014). In general, following the original KcsA crystal structure, which indicated alternating water and K^+^ ions within the sequential SF binding sites, these studies have assumed that conduction involves concerted movement of ions and water through the filter, requiring combined electrical and steric forces(Åqvist & Luzhkov, 2000),(Berneche & Roux, 2001). It has also been suggested that less coordinated transitions may be involved(Åqvist & Luzhkov, 2000; Furini & Domene, 2009), and more recent studies have suggested that, rather than alternating ions and water, K^+^ ions can occupy adjacent sites in the SF and conductance is then governed by a direct knock-on mechanism(Kopec et al, 2018; Köpfer et al, 2014; Langan et al, 2018), such as we observe here for Kir2.2. Except for the starting configuration, K^+^ ions unambiguously permeated the open Kir2.2 channel in a fully dehydrated state, via a direct knock-on mechanism, in which an ion entering the S4 site pushed the ion ahead to site S3 and then S2, leading to exit of the outermost ion on the extracellular side. While a direct knock-on mechanism is thus a clear finding here, one potential caveat is that certain features of the SF may not be fully accounted for in MD. The Tyr side chain in the SF loop has to be constrained to maintain a conductive SF conformation in the simulations, otherwise, as noted by others, flipping of peptide bonds can result in a non-conducting distortion(Domene et al, 2004; Haider et al, 2007). Multiple experimental studies indicate that subtle conformational changes or mutations at this residue have significant effects on permeation. Conservative mutation from Tyr (Y) to Phe (Y145F) in KcsA, as well as to other residues(Liu et al, 2001) affects apparent channel open times, while in K_ATP_ (Kir6.2) channels, conservative mutation of the equivalent Phe at residue 133 to Tyr results in non-functional channels(Proks et al, 2001). Cryo-EM structures of WT and inactivation-deficient S631A mutant hERG (Kv11.1) channels (Schonherr & Heinemann, 1996; Wang & MacKinnon, 2017) also suggest subtle conformational changes of the aromatic side chain position of the equivalent F627 residue could underlie channel inactivation.

It has long been recognized that water molecules traverse K^+^ channels as well as the permeant ion(Alcayaga et al, 1989). Intriguingly, the coupling ratio between water and K^+^ typically declines in more K^+^-selective channels(Kopec et al, 2018), and as [K^+^] increases(Ando et al, 2005), consistent with water movement being reduced as K^+^ occupancy increases. Our recent single molecule FRET studies indicate that the canonical K^+^-selective conformation of the KirBac1.1 SF is stabilized by K^+^, and that significant filter dilation, resulting in a non-selective conformation, occurs at lower [K^+^](Wang et al, 2019). While it is unknown how generalizable these findings are, it raises the notion that the apparent coupling of water and K^+^ ion flux could be a result of the SF cycling between two distinct conformations, one in which water-independent K^+^ permeation occurs through the canonical SF conformation via a direct K^+^-ion knock on mechanism, and one in which water permeates through the dilated SF conformation.

## Conclusions

We have generated mutant eukaryotic Kir2.2 channels in which both known lipid-gating requirements are met and that force channel opening in recombinant channels. Despite this, crystal structures remained in a non-conducting state. However, in subsequent MD simulations, in which crystallographic constraints are removed, the mutant channel pores spontaneously widened at the HBC gate, resulting in ‘wetting’ at the HBC region, high K^+^ concentrations in the inner cavity, and K^+^ ion permeation at comparable rates to measured ion conductance. When channels were ‘back mutated’ to the wild type Gly at residue 178 after ‘wetting’ had occurred, channels remained in an open conductive state, but with intermittent ‘de-wetted’ phases and reduced overall conductance, paralleling lower experimentally measured single channel conductance in the WT versus mutant channel. Ion conduction through the pore occurred strictly via a direct knock-on mechanism, as observed in simulations of other potassium channels. The results provide key insights to gating and permeation of inward rectifier K channels.

## Materials and Methods

### Cloning, expression and purification

A single point mutation (G178D) was introduced using QuickChange site-directed mutagenesis (Stratagene Cloning Systems, CA) to both the wild type and K62W mutant truncated cKir2.2 channel cDNA and verified by sequencing. Mutant channel proteins were expressed in *Pichia Pastoris* cells as described previously(Tao et al, 2009). Briefly, frozen *Pichia* cells were broken using a Retsch, Inc. Model MM400 mixer mill (5×3.0 minutes at 25 CPS) and solubilized in lysis buffer (100 mM DM, 50 mM Tris pH 7.5, 150 mM KCl, 1 mM EDTA, 2 mM DTT) at room temperature for 1 hour with stirring or rotation. Cell lysate was centrifuged at 30,000g for 30 min at 10 °C. Approximately, 750 μL of Anti-Flag Ab conjugated resins per 10 g of cells was added to the supernatant. Binding was performed at 4 °C for >1.5 hour with gentle rotation. The resin was washed with 20 column volumes of wash buffer (4 mM DM, 50 mM Tris pH 7.5, 150 mM KCl, 1 mM EDTA, 2 mM DTT), and protein was released from the resin by PreScission Protease (PPX) action at 4 °C overnight and further purified on a Superdex 200 column (GE Healthcare) equilibrated in SEC buffer (20 mM Tris pH 7.5, 150 mM KCl, 1 mM EDTA, 4 mM DM, 20 mM DTT). Peak fractions were combined and concentrated to >5 mg/ml for crystallization experiments and *in vitro* functional assays.

### Electrophysiology

CosM6 cells were transfected with 1-2 µg of pcDNA3.1-Kir2.2-K62W or pcDNA3.1-Kir2.2-K62W-G178D mutants with an addition of 0.4 µg of pcDNA3.1-GFP per 35mm Petri dish using FuGENE6 (Promega). The cells were used for patching within 1-2 days after transfection. For patch-clamp experiments, symmetrical internal potassium buffer (K_int_) was used: 148 mM KCl, 1 mM EGTA, 1 mM K_2_EDTA, 10 mM HEPES (pH 7.4). Data were acquired at 15 kHz, low-pass filtered at 5 kHz with Axopatch 1D patch-clamp amplifier and digitized with Digidata 1320 digitizer (Molecular Devices). Data analysis was performed using the pClamp software suite (Molecular Devices). Pipettes with bubble number (BN)(Schnorf et al, 1994) 4.0 – 5.0 (~ 2-4 MOhm in symmetric K_int_) were pulled from Kimble Chase 2502 soda lime glass with a Sutter P-86 puller (Sutter Instruments). All measurements were carried out on excised inside-out patches at-120 mV membrane potential, or as specified in the text.

### Rb^+^ Flux Assay

Channel activity of purified recombinant proteins was assessed by _86_Rb^+^ uptake into proteoliposomes. 1-palmitoyl-2-oleoyl-sn-glycero-3-phosphoethanolamine (POPE) and 1-palmitoyl-2-oleoyl-sn-glycero-3-phospho-(1’-rac-glycerol) (POPG) lipids were dissolved (10 mg/ml) in buffer A (450 mM KCl, 10 mM HEPES, 4 mM NMG, 0.5 mM EGTA, pH 7.4) with 35 mM CHAPS. Porcine brain PIP_2_ was solubilized (2 mg/ml) in 8 mg/ml POPE solution. 1 mg of lipid mixture in 100 µl was incubated at room temperature for 2 hours, and 3 – 10 µg of protein was added and incubated for an additional 20 mins. The lipid-protein mixture was passed through partly dehydrated G-50 beads pre-equilibrated with buffer A to form proteoliposomes. Proteoliposomes were passed through partly dehydrated G-50 beads pre-equilibrated by buffer B (400 mM Sorbitol, 10 mM Hepes, 4 mM NMDG, 0.5 mM EGTA, pH 7.4), to remove external KCl. For assessing ion selectivity, the internal salt was replaced with Li, Na, Rb, Cs, and NMDG. For blocking experiments, spermine was added only to the external side to a concentration of 10 μM; higher concentrations could not be assessed because of liposome destabilization (data not shown). 200 µl of buffer B containing _86_Rb^+^ (PerkinElmer, Waltham, MA) at approximately 4.5 nCi was added at time zero. After 10 mins, 80 µl samples were collected and passed through cation exchange beads to capture ^86^Rb^+^ in the external solution. Uptake was normalized to the maximum uptake in valinomycin, after leak subtraction (uptake into protein-free liposomes). Porcine brain PIP_2_, POPE, and POPG were purchased from Avanti Polar Lipids.

### Crystallization and structure determination

1,2-dioleoyl-sn-glycero-3-phospho-(1’-myo-inositol-4’,5’-bisphosphate) (PIP_2_) was solubilized in SEC buffer (5 mM) and added to the concentrated protein sample to 250 μM concentration, at least 30 min prior to crystallization. Crystals were grown by hanging drop vapor diffusion, by mixing 0.5 µL of protein and 0.5 µL of reservoir solutions. Crystals for the Apo-KW/GD structure were obtained in 180 mM triNaCitrate, 100 mM Tris pH 7.1, and 27.4% PEG 400 at 20 °C; PIP_2_-KW/GD crystals were obtained in 80 mM triNaCitrate, 100 mM Tris pH 7.1, and 27.2% PEG 400 at 20 °C. Crystals grew at 20 °C in 2-3 days and were cryoprotected by 30 % (v/v) PEG400 and frozen in liquid nitrogen. X-ray diffraction data (3.6 Å for Apo, 2.81 Å for PIP_2_-crystals) were collected at the Advanced Photon Source beamlines 24-ID-C and 24-ID-E at wavelength 0.9792 Å under liquid nitrogen stream. Phases were obtained using the program MolRep(Vagin & Teplyakov, 2000) in the CCP4 suite(Winn et al, 2011) through molecular replacement with Apo-K62W mutant crystal structure (PDB code 5KUM) as a search model for PIP_2_-KW/GD crystals, and 5SPC (cKir2.2 WT co-crystallized with pyrophosphatidic acid (PPA)) for Apo-KW/GD crystals. Iterative model building was carried out in COOT(Emsley & Cowtan, 2004), and rounds of refinement were performed with REFMAC(Murshudov et al, 1997). The final Apo-KW/GD model (PDB ID: 6M86) was obtained with R/R_free_ of 0.237/0.291 and with 96% of residues in the most favored regions and one in the disallowed region in the Ramachandran plot. The final PIP_2_-KW/GD model (PDB ID: 6M84) was obtained with R/R_free_ of 0.22/0.270 and with 97% of residues in the most favored regions and none in the disallowed region in the Ramachandran plot.

### Molecular dynamics simulations

MD simulations and subsequent analyses were performed using the software package Gromacs v.5.1.2(Abraham et al, 2015). As a starting point, the structure was inserted into an equilibrated bilayer membrane consisting of palmitoyloleoylphosphatidylcholine (POPC) lipids, for which Berger lipid parameters were employed(Berger et al, 1997; Cordomí et al, 2012), surrounded by explicit water molecules (SPCE water model(Berendsen et al, 1987)) and 0.2 M potassium chloride(Joung & Cheatham, 2008). Geometry optimization and electrostatic potential calculation of PIP_2_ (short chain analogue) were performed at the Hartree-Fock/6-31G* level using the program Gaussian09 (Gaussian, Inc., Wallingford, CT, USA). The generalized amber force field topologies of the ligand were generated with the Antechamber software(Wang et al, 2006) using partial charges from quantum mechanics calculations according to the restrained electrostatic potential (RESP) approach. Potassium ions were placed in the selectivity filter at sites S0, S2, and S4, and water molecules were placed at sites S1 and S3(Åqvist & Luzhkov, 2000). The amber99sb force field was used for the protein(Hornak et al, 2006). For the ions we used corrected monovalent Lennard-Jones parameters for the amber force field(Joung & Cheatham, 2008). The cut-off for electrostatic interactions was set to 1.0 nm; long-range electrostatic interactions were treated by the particle-mesh Ewald method(Darden et al, 1993). The cut-off for Lennard-Jones interactions was set to 1.0 nm. The LINCS algorithm was used to constrain bonds(Hess et al, 1997). The simulation temperature was kept constant using a V-rescale thermostat(Bussi et al, 2007); protein and lipids as well as the solvent, together with ions were coupled (τ = 0.1 ps) separately to a temperature bath of 310 K. Similarly, the pressure was kept constant at 1 bar by the Parrinello-Rahman barostat algorithm (coupling constant = 2 ps)(Bussi et al, 2007). Steepest descent energy-minimization was performed prior to all simulations. To equilibrate the system, we position-restrained all heavy atoms with a force constant (fc) of 1000 kJmol^−1^nm^−2^ and simulated for 20 ns. The Y146 side chain atoms were restrained throughout the simulations with a fc of 1000 kJmol^−1^nm^−2^. An electric field along the channel pore (40 mV·nm^−1^) was applied. With a box length of ~14.6 nm, this amounts to a transmembrane potential of ~580 mV. *In silico* mutation of G178D-> G (G178D(G)) was accomplished using Swiss PDB Viewer(Guex & Peitsch, 1997).

### Analysis of MD trajectories

Changes in the relative rotation of the CTD with respect to the TMD were assessed by calculating the torsion angle between two planes. Calculation of the torsion angle required four different points of measurement: the center of mass of the TMD (point 1), the center of mass of the CTD (point 2), the center of mass of one subunit of the TMD (point 3) and the center of mass of one subunit of the CTD (point 4). The first plane is defined by points 1, 2 and 3; the second plane is defined by points 1, 2 and 4. To calculate the changes over simulation time, the angle between these two planes of the first simulation step was defined as zero (for similar methods see Linder et al.(Linder et al, 2015)).

Before analyzing water occupancies/solvation, all trajectories were aligned at the selectivity filter (sequence TTIGYG). The coordinates of protein and potassium ions were saved every 20ps. Depending on simulation length, this resulted in 10,000 steps (for 200ns trajectories) or 50,000 steps (for 1µs trajectories) per run. Along the membrane normal (= pore axis z), the area between the intracellular entrance of the channel and the end of the SF was cut into slices of 0.5 Å thickness. Potassium ions inside the channel pore were counted in each slice. Average numbers of resident potassium ions were plotted against the membrane normal. Based on these occupancies the PMF (potential of mean force) was determined using the following equation: G_PMF_(z) = −kBT ln n(z)(de Groot & Grubmüller, 2001).

### Statistical Analysis

Statistical significance was analyzed using unpaired T-tests unless otherwise stated. Statistical significance of *p*<0.05, *p*<0.01, and *p*<0.001 is indicated by single, double, and triple asterisks respectively.

## Supporting information

Movie 1 (Quicktime)

Movie 4 (Quicktime)

Movie 2 (Quicktime)

Movie 3 (Quicktime)

## Acknowledgements

We thank the staffs at APS beamlines 23-ID-C/E, especially K. Rajashankar, K. Perry and N. Sukumar, for assistance at the synchrotron. This work used NE-CAT beamlines (GM103403), a Pilatus detector (RR029205) and an Eiger detector (OD021527) at the APS (DE-AC02-06CH11357). This work was supported by NIH grant HL140024 (to CGN), and Austrian Science Fund grants W1232 (to E-MZP, HB and ASW) and I-2101-B26 (to ASW). The computational results presented have been achieved using the Vienna Scientific Cluster (VSC) and XSEDE. We thank Peter Zangerl for computational advice and scripting.

## Author contributions

E-MZP, SJL, ASW and CGN conceived the project. SJL and GM carried out experiments. E-MZP and HB carried out molecular modeling and analyses. FR and PY collected the diffraction data, which was analyzed by SJL. E-MZP, SJL, ASW, and CGN wrote the paper, which was edited by HB, GM, FR and PY.

## Data availability

The datasets and computer code produced in this study are available upon request, and will be deposited in appropriate public database.

## References

Abraham MJ, Murtola T, Schulz R, Páll S, Smith JC, Hess B, Lindahl E (2015) GROMACS: High performance molecular simulations through multi-level parallelism from laptops to supercomputers. SoftwareX 1-2: 19–25

Alcayaga C, Cecchi X, Alvarez O, Latorre R (1989) Streaming potential measurements in Ca2+-activated K+ channels from skeletal and smooth muscle. Coupling of ion and water fluxes. Biophys J 55: 367–371

Ando H, Kuno M, Shimizu H, Muramatsu I, Oiki S (2005) Coupled K+-water flux through the HERG potassium channel measured by an osmotic pulse method. J Gen Physiol 126: 529–538

Åqvist J, Luzhkov V (2000) Ion permeation mechanism of the potassium channel. Nature 404: 881–884

Aryal P, Sansom MS, Tucker SJ (2015) Hydrophobic gating in ion channels. J Mol Biol 427: 121–130

Bavro VN, De Zorzi R, Schmidt MR, Muniz JR, Zubcevic L, Sansom MS, Venien-Bryan C, Tucker SJ (2012) Structure of a KirBac potassium channel with an open bundle crossing indicates a mechanism of channel gating. Nat Struct Mol Biol 19: 158–163

Berendsen H, Grigera J, Straatsma T (1987) The missing term in effective pair potentials. Journal of Physical Chemistry 91: 6269–6271

Berger O, Edholm O, Jähnig F (1997) Molecular dynamics simulations of a fluid bilayer of dipalmitoylphosphatidylcholine at full hydration, constant pressure, and constant temperature. Biophysical Journal 72: 2002–2013

Berneche S, Roux B (2001) Energetics of ion conduction through the K+ channel. Nature 414: 73–77

Bradley J, O’Meara F, Farrell D, Nielsen JE (2012) Highly perturbed pKa values in the unfolded state of hen egg white lysozyme. Biophys J 102: 1636–1645

Brelidze TI, Niu X, Magleby KL (2003) A ring of eight conserved negatively charged amino acids doubles the conductance of BK channels and prevents inward rectification. Proc Natl Acad Sci U S A 100: 9017–9022

Bussi G, Donadio D, Parrinello M (2007) Canonical sampling through velocity rescaling. The Journal of Chemical Physics 126: 014101

Cheng WWL, D’Avanzo N, Doyle DA, Nichols CG (2011) Dual-Mode Phospholipid Regulation of Human Inward Rectifying Potassium Channels. Biophysical Journal 100: 620–628

Cordomí A, Caltabiano G, Pardo L (2012) Membrane Protein Simulations Using AMBER Force Field and Berger Lipid Parameters. Journal of Chemical Theory and Computation 8: 948–958

D’Avanzo N, Cheng WW, Doyle DA, Nichols CG (2010) Direct and specific activation of human inward rectifier K+ channels by membrane phosphatidylinositol 4,5-bisphosphate. J Biol Chem 285: 37129–37132

Dahl AC, Chavent M, Sansom MS (2012) Bendix: intuitive helix geometry analysis and abstraction. Bioinformatics 28: 2193–2194

Darden T, York D, Pedersen L (1993) Particle mesh Ewald: An N⋅ log (N) method for Ewald sums in large systems. The Journal of chemical physics 98: 10089–10092

de Groot BL, Grubmüller H (2001) Water permeation across biological membranes: mechanism and dynamics of aquaporin-1 and GlpF. Science 294: 2353–2357

Diaz-Franulic I, Sepulveda RV, Navarro-Quezada N, Gonzalez-Nilo F, Naranjo D (2015) Pore dimensions and the role of occupancy in unitary conductance of Shaker K channels. J Gen Physiol 146: 133–146

Domene C, Grottesi A, Sansom MS (2004) Filter flexibility and distortion in a bacterial inward rectifier K+ channel: simulation studies of KirBac1.1. Biophys J 87: 256–267

Emsley P, Cowtan K (2004) Coot: model-building tools for molecular graphics. Acta Crystallogr D Biol Crystallogr 60: 2126–2132

Fujiwara Y, Kubo Y (2006) Functional roles of charged amino acid residues on the wall of the cytoplasmic pore of Kir2.1. J Gen Physiol 127: 401–419

Furini S, Domene C (2009) Atypical mechanism of conduction in potassium channels. Proc Natl Acad Sci U S A 106: 16074–16077

Gosink LJ, Hogan EA, Pulsipher TC, Baker NA (2014) Bayesian model aggregation for ensemble-based estimates of protein pKa values. Proteins 82: 354–363

Grandi E, Sanguinetti MC, Bartos DC, Bers DM, Chen-Izu Y, Chiamvimonvat N, Colecraft HM, Delisle BP, Heijman J, Navedo MF, Noskov S, Proenza C, Vandenberg JI, Yarov-Yarovoy V (2017) Potassium channels in the heart: structure, function and regulation. J Physiol 595: 2209–2228

Guex N, Peitsch MC (1997) SWISS-MODEL and the Swiss-Pdb Viewer: An environment for comparative protein modeling. ELECTROPHORESIS 18: 2714–2723

Haider S, Khalid S, Tucker SJ, Ashcroft FM, Sansom MS (2007) Molecular dynamics simulations of inwardly rectifying (Kir) potassium channels: a comparative study. Biochemistry 46: 3643–3652

Hansen SB, Tao X, MacKinnon R (2011) Structural basis of PIP2 activation of the classical inward rectifier K+ channel Kir2.2. Nature 477: 495–498

Hess B, Bekker H, Berendsen HJC, Fraaije JGEM (1997) LINCS: A linear constraint solver for molecular simulations. Journal of Computational Chemistry 18: 1463–1472

Hibino H, Inanobe A, Furutani K, Murakami S, Findlay I, Kurachi Y (2010) Inwardly Rectifying Potassium Channels: Their Structure, Function, and Physiological Roles. Physiological Reviews 90: 291–366

Hilgemann DW, Ball R (1996) Regulation of cardiac Na+,Ca2+ exchange and KATP potassium channels by PIP2. Science 273: 956–959

Hite RK, MacKinnon R (2017) Structural Titration of Slo2.2, a Na(+)-Dependent K(+) Channel. Cell 168: 390–399 e311

Ho BK, Gruswitz F (2008) HOLLOW: generating accurate representations of channel and interior surfaces in molecular structures. BMC Struct Biol 8: 49

Hornak V, Abel R, Okur A, Strockbine B, Roitberg A, Simmerling C (2006) Comparison of multiple Amber force fields and development of improved protein backbone parameters. Proteins: Structure, Function, and Bioinformatics 65: 712–725

Joung IS, Cheatham TE (2008) Determination of Alkali and Halide Monovalent Ion Parameters for Use in Explicitly Solvated Biomolecular Simulations. The Journal of Physical Chemistry B 112: 9020–9041

Kopec W, Kopfer DA, Vickery ON, Bondarenko AS, Jansen TLC, de Groot BL, Zachariae U (2018) Direct knock-on of desolvated ions governs strict ion selectivity in K(+) channels. Nat Chem 10: 813–820

Köpfer DA, Song C, Gruene T, Sheldrick GM, Zachariae U, de Groot BL (2014) Ion permeation in K^+^ channels occurs by direct Coulomb knock-on. Science 346: 352–355

Kurata HT, Zhu EA, Nichols CG (2010) Locale and chemistry of spermine binding in the archetypal inward rectifier Kir2.1. J Gen Physiol 135: 495–508

Langan PS, Vandavasi VG, Weiss KL, Afonine PV, El Omari K, Duman R, Wagner A, Coates L (2018) Anomalous X-ray diffraction studies of ion transport in K(+) channels. Nat Commun 9: 4540

Lee S-J, Wang S, Borschel W, Heyman S, Gyore J, Nichols CG (2013) Secondary anionic phospholipid binding site and gating mechanism in Kir2.1 inward rectifier channels. Nat Commun 4

Lee SJ, Ren F, Zangerl-Plessl EM, Heyman S, Stary-Weinzinger A, Yuan P, Nichols CG (2016) Structural basis of control of inward rectifier Kir2 channel gating by bulk anionic phospholipids. J Gen Physiol 148: 227–237

Li J, Lu S, Liu Y, Pang C, Chen Y, Zhang S, Yu H, Long M, Zhang H, Logothetis DE, Zhan Y, An H (2015) Identification of the Conformational transition pathway in PIP2 Opening Kir Channels. Sci Rep 5: 11289

Linder T, Wang S, Zangerl-Plessl E-M, Nichols CG, Stary-Weinzinger A (2015) Molecular Dynamics Simulations of KirBac1.1 Mutants Reveal Global Gating Changes of Kir Channels. Journal of Chemical Information and Modeling 55: 814–822

Liu YS, Sompornpisut P, Perozo E (2001) Structure of the KcsA channel intracellular gate in the open state. Nat Struct Biol 8: 883–887

Lopatin AN, Makhina EN, Nichols CG (1994) Potassium channel block by cytoplasmic polyamines as the mechanism of intrinsic rectification. Nature 372: 366–369

Martin GM, Yoshioka C, Rex EA, Fay JF, Xie Q, Whorton MR, Chen JZ, Shyng SL (2017) Cryo-EM structure of the ATP-sensitive potassium channel illuminates mechanisms of assembly and gating. Elife 6

Murshudov GN, Vagin AA, Dodson EJ (1997) Refinement of macromolecular structures by the maximum-likelihood method. Acta Crystallogr D Biol Crystallogr 53: 240–255

Naranjo D, Moldenhauer H, Pincuntureo M, Diaz-Franulic I (2016) Pore size matters for potassium channel conductance. J Gen Physiol 148: 277–291

Nichols CG, Lopatin AN (1997) Inward rectifier potassium channels. Annu Rev Physiol 59: 171–191

Nimigean CM, Chappie JS, Miller C (2003) Electrostatic tuning of ion conductance in potassium channels. Biochemistry 42: 9263–9268

Phillips LR, Nichols CG (2003) Ligand-induced closure of inward rectifier Kir6.2 channels traps spermine in the pore. J Gen Physiol 122: 795–804

Proks P, Capener CE, Jones P, Ashcroft FM (2001) Mutations within the P-loop of Kir6.2 modulate the intraburst kinetics of the ATP-sensitive potassium channel. J Gen Physiol 118: 341–353

Schnorf M, Potrykus I, Neuhaus G (1994) Microinjection technique: routine system for characterization of microcapillaries by bubble pressure measurement. Exp Cell Res 210: 260–267

Schonherr R, Heinemann SH (1996) Molecular determinants for activation and inactivation of HERG, a human inward rectifier potassium channel. J Physiol 493 (Pt 3): 635–642

Smart OS, Neduvelil JG, Wang X, Wallace BA, Sansom MS (1996) HOLE: a program for the analysis of the pore dimensions of ion channel structural models. J Mol Graph 14: 354–360, 376

Takahashi N, Morishige K, Jahangir A, Yamada M, Findlay I, Koyama H, Kurachi Y (1994) Molecular cloning and functional expression of cDNA encoding a second class of inward rectifier potassium channels in the mouse brain. J Biol Chem 269: 23274–23279

Tao X, Avalos JL, Chen J, MacKinnon R (2009) Crystal structure of the eukaryotic strong inward-rectifier K+ channel Kir2.2 at 3.1 A resolution. Science 326: 1668–1674

Trick JL, Aryal P, Tucker SJ, Sansom MS (2015) Molecular simulation studies of hydrophobic gating in nanopores and ion channels. Biochem Soc Trans 43: 146–150

Vagin A, Teplyakov A (2000) An approach to multi-copy search in molecular replacement. Acta Crystallogr D Biol Crystallogr 56: 1622–1624

Wang J, Wang W, Kollman PA, Case DA (2006) Automatic atom type and bond type perception in molecular mechanical calculations. Journal of molecular graphics & modelling 25: 247–260

Wang S, Lee S-JL, Maksiev G, Fang X, Zuo C, Nichols CG (2019) Potassium channel selectivity filter dynamics revealed by single-molecule FRET. Nature Chemical Biology (In press)

Wang S, Lee SJ, Heyman S, Enkvetchakul D, Nichols CG (2012) Structural rearrangements underlying ligand-gating in Kir channels. Nat Commun 3: 617

Wang S, Vafabakhsh R, Borschel WF, Ha T, Nichols CG (2016) Structural dynamics of potassium-channel gating revealed by single-molecule FRET. Nat Struct Mol Biol 23: 31–36

Wang W, MacKinnon R (2017) Cryo-EM Structure of the Open Human Ether-a-go-go-Related K(+) Channel hERG. Cell 169: 422–430 e410

Whorton Matthew R, MacKinnon R (2011) Crystal Structure of the Mammalian GIRK2 K+ Channel and Gating Regulation by G Proteins, PIP2, and Sodium. Cell 147: 199–208

Whorton MR, MacKinnon R (2013) X-ray structure of the mammalian GIRK2-[bgr][ggr] G-protein complex. Nature 498: 190–197

Winn MD, Ballard CC, Cowtan KD, Dodson EJ, Emsley P, Evans PR, Keegan RM, Krissinel EB, Leslie AG, McCoy A, McNicholas SJ, Murshudov GN, Pannu NS, Potterton EA, Powell HR, Read RJ, Vagin A, Wilson KS (2011) Overview of the CCP4 suite and current developments. Acta Crystallogr D Biol Crystallogr 67: 235–242

Xu Y, Shin HG, Szep S, Lu Z (2009) Physical determinants of strong voltage sensitivity of K(+) channel block. Nat Struct Mol Biol 16: 1252–1258

