## Supplementary Data for "Atomistic basis of opening and conduction in mammalian inward rectifier potassium (Kir2.2) channels"

**Supplementary Table 1: Crystallographic data and refinement statistics**

| | Apo-KWGD (6M86) | $PIP_2$-bound KWGD (6M84) |
|---|---|---|
| **Data collection** | | |
| Space group | I4 | I4 |
| Cell dimensions | | |
| a, b, c (Å) | 82.852, 82.852, 189.109 | 82.878 82.787 182.976 |
| $\alpha = \beta = \gamma$ (°) | 90, 90, 90 | 90, 90, 90 |
| Resolution (Å) | 50 – 3.6 (3.73 – 3.60) | 91.95 – 2.80 (2.90 – 2.80) |
| Unique Reflections | 7432 | 15066 |
| $R_{merge}^{*}$ | 0.079 (0.742) | 0.059 (0.947) |
| $I / \sigma I$ | 273.6/9.7 (5.9/2.1) = 29.1 (2.8) | 168.4/5.6 (4.3/2.0) = 30.1 (2.2) |
| Completeness (%) | 99.9(100.0) | 99.9 (99.9) |
| Redundancy | 3.4 (3.2) | 3.4 (3.3) |
| **Refinement** | | |
| Resolution (Å) | 50 – 3.6 | 50 – 2.80 |
| No. of reflections | 7040 | 14242 |
| $R_{work}/R_{free}$ | 0.237/0.291 | 0.222/0.270 |
| No. atoms | | |
| Protein | 2578 | 2620 |
| Ligand/ion | 0/5 | 52/6 |
| Water | 5 | 19 |
| B-factors | | |
| Protein | 187.0 | 98.5 |
| Ligand/ion | 0/151.3 | 164.3/101.5 |
| Water | 138.704 | 81.7 |
| R.m.s. deviation | | |
| Bond length (Å) | 0.004 | 0.005 |
| Bond angles (°) | 0.851 | 1.027 |

$^{*}$ $R_{merge}$ = SUM(| I -<I>|)/SUM(I)

**Supplementary Table 2: Overview of performed simulations.**

| Simulation setup | Number of runs | Simulation time [ns] /run | Sum of Simulation time [ns] | Number of ions passed through the SF | Unitary conductance (pS) |
|---|---|---|---|---|---|
| KWGD restrained HBC gate | 1 | 500 | 500 | 0 | 0 |
| KWGD(G) start | 1 | 100 | 100 | 0 | 0 |
| KWGD | 4 | 200 | 2,800 | 6,4,1,0 | 8.43, 5.62, 1.41, 0 |
| | 2 | 1,000 | | 14, 6 | 3.94, 1.69 |
| KWGD(G) after 50ns | 10 | 200 | 3,000 | 4, 1, 1, 0, 0, 0, 0, 0, 0, 0 | 5.62, 1.41, 1.41, 0, 0, 0, 0, 0, 0, 0 |
| | 1 | 1000 | | 6 | 1.69 |

**Supplementary Table 3: predicted pKa values of ionizable residues along the pore axis by APBS**

| Residue | $PIP_2$_KW (xtal:closed) | $PIP_2$-KW/GD (xtal:closed) | $PIP_2$-KW/GD (MD:open) | $PIP_2$-KW/GD(G) (MD:open) |
|---|---|---|---|---|
| **D173** | 7.04<br>7.32<br>7.04<br>7.32 | 7.12<br>7.21<br>7.12<br>7.21 | 6.50<br>5.83<br>6.50<br>5.88 | 5.44<br>5.90<br>6.77<br>5.87 |
| **G178D** | - | 9.92<br>9.39<br>9.92<br>9.39 | 5.99<br>5.77<br>6.38<br>5.43 | - |
| **E225** | 5.72<br>6.09<br>5.72<br>6.09 | 3.32<br>5.89<br>3.32<br>5.89 | 9.35<br>7.21<br>5.09<br>7.75 | 8.81<br>6.24<br>2.54<br>6.85 |
| **E300** | 2.11<br>2.11<br>2.11<br>2.11 | 3.54<br>5.69<br>3.54<br>5.69 | 5.92<br>0.67<br>3.92<br>3.09 | 3.45<br>6.50<br>2.64<br>6.32 |
| **D256** | 3.95<br>4.01<br>3.95<br>3.92 | 4.00<br>3.87<br>4.00<br>3.87 | 4.30<br>3.98<br>4.16<br>2.69 | 4.38<br>4.01<br>4.17<br>2.82 |

**Supplementary Figures**

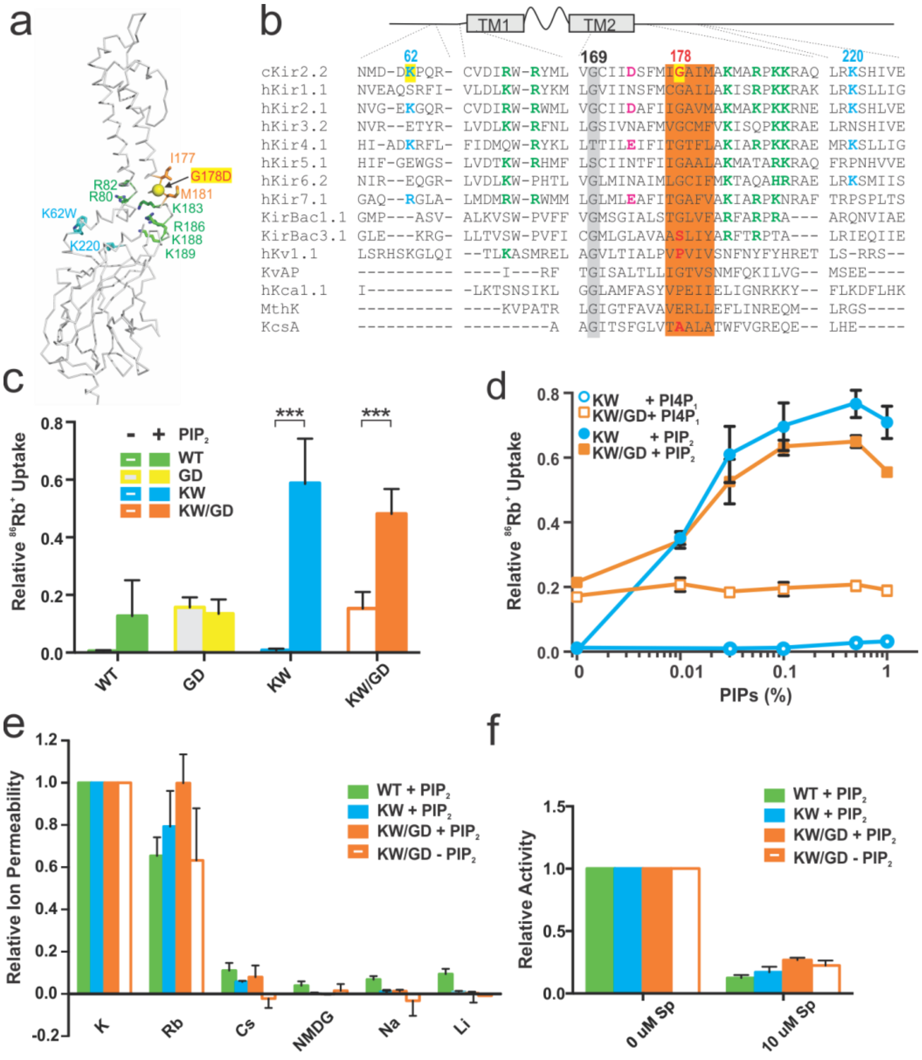


**Supplementary Figure S1. Force open mutant channel properties *in vitro*.** a) Backbone trace of Kir2 channel with the side chain of the two residues (I177 and M181) forming the helix bundle crossing (HBC) in sticks and the mutated residue G178 in balls. $PIP_2$ binding site residues are shown in sticks and green and bulk anionic lipid binding site residues are in sticks and cyan. B) Sequence alignment of $K^+$ channels. The residues at the $PIP_2$ binding site are shown in green and the residues important for secondary anionic lipids in cyan. The residues are mutated in this study are designated by yellow box, and the residues considered to form HBC are marked by an orange shade. Negatively charged residue in the inner cavity important for rectification, so called ‘rectification controller’, are shown in pink and the conserved glycine hinge residues are shaded in gray. c) Mutant and control protein activity was assessed in the absence and presence of 0.1% $PIP_2$ in the liposomes made with 10% POPG and 90% POPE. d) $PIP_2$ specificity was tested for KW and KW/GD proteins. $PIP_2$ or $PI(4)P_1$ were increased from zero to 1% in the liposomes made of 10% POPG and 90% POPE. e) Ion selectivity of the proteins was assessed by generating concentration gradient with different ions inside the proteoliposomes while the lipid composition was kept the same (0.1% $PIP_2$, 10% POPG, and 90% POPE). f) Pore blocking by spermine was assessed through $Rb^+$ flux assay by adding 10 µM spermine to the external buffer. The lipid composition was kept the same (0.1% $PIP_2$, 10% POPG, and 90% POPE). Columns and symbols are means, and error bars are standard errors (n >= 3).

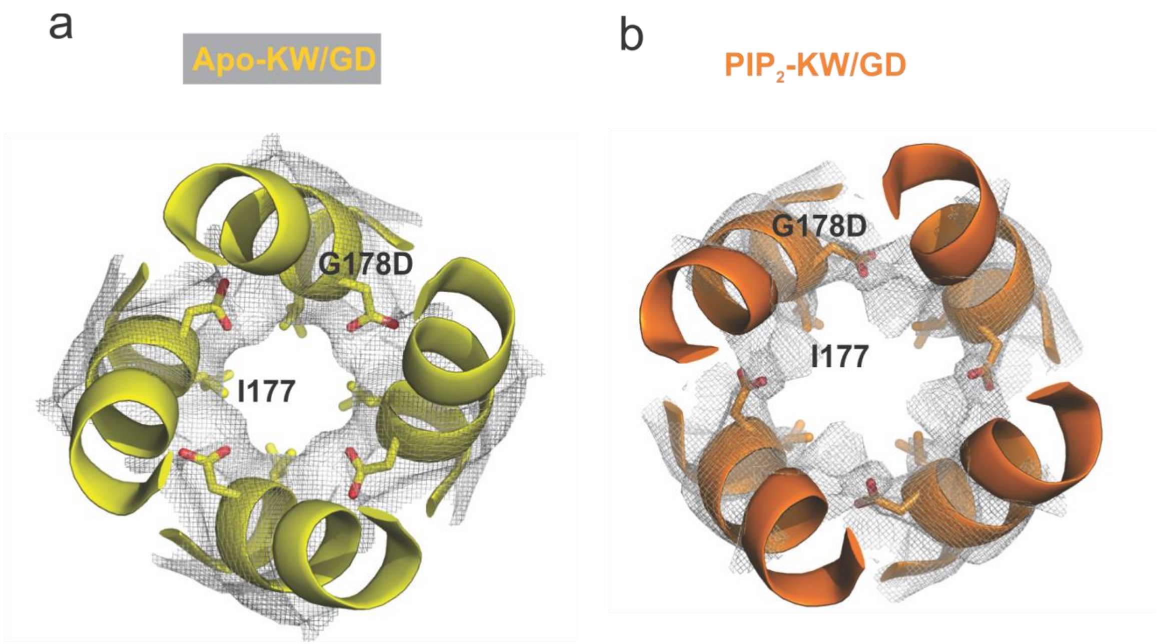


**Supplementary Figure S2. G178D residues in crystal structures.** Bottom-up views (in the plane of the membrane) of the two crystal structures near the HBC are shown in ribbon diagrams with electron densities contoured at 1 σ. The mutant G178D and HBC forming I177 residues are drawn explicitly in sticks.

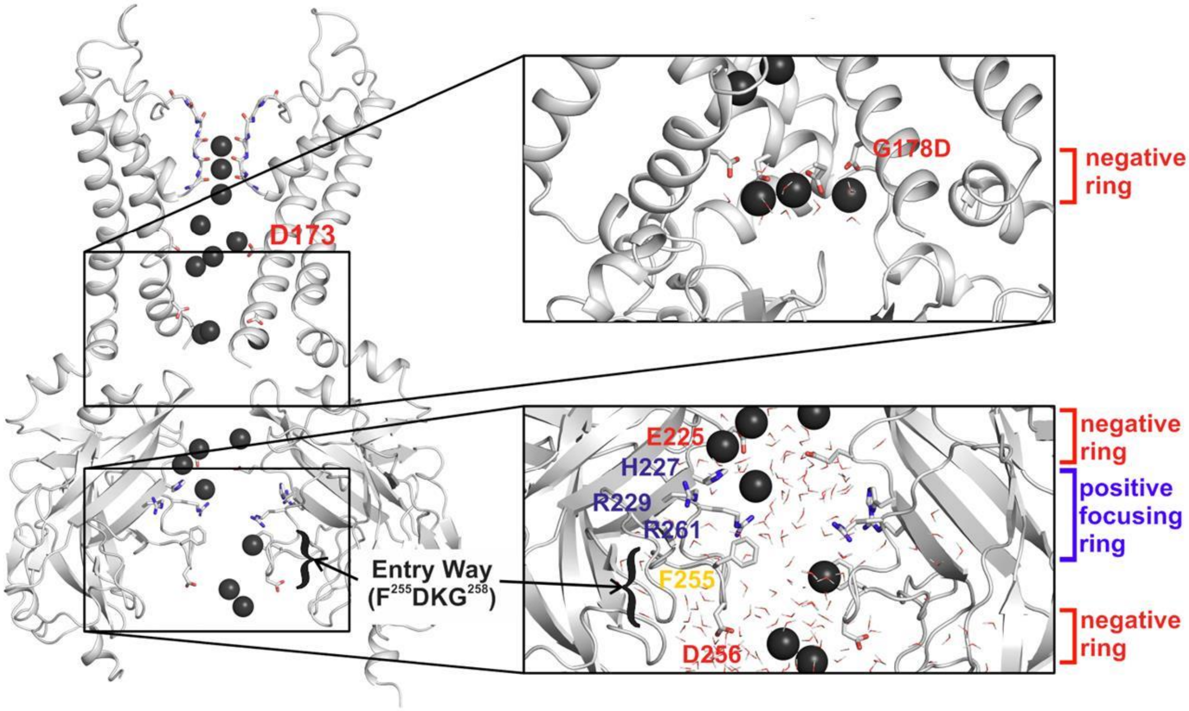


**Supplementary Figure S3. A snapshot of $K^+$ ion distribution along the open pore during $PIP_2$-KW/GD MD simulation.** Two opposing subunits are shown in ribbons and $K^+$ ions and water molecules in the pore are shown in black balls and sticks respectively. Residues of interest are shown in sticks and labeled. Zoom-in views show the rings of acidic (negative) and basic (positive) residues along the pore.

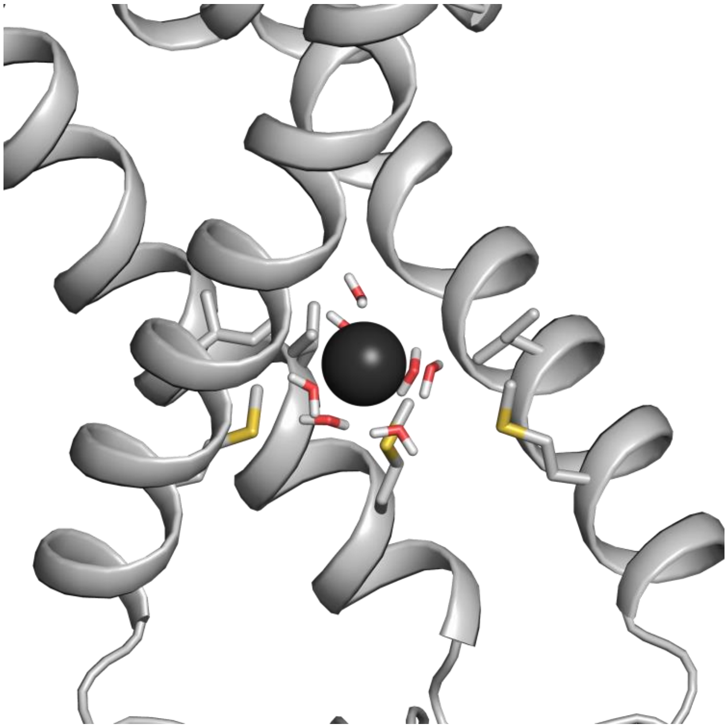

**Supplementary Figure S4. Snapshot of a solvated ion passing the HBC gate.** TM2 helices are represented as gray cartoon, the subunit in the front is omitted for clarity. Residues I177 and M181 as well as water molecules are shown in sticks. The $K^+$ ion at the HBC is represented as a black sphere. This snapshot was taken from a $PIP_2$-KW/GD simulation.

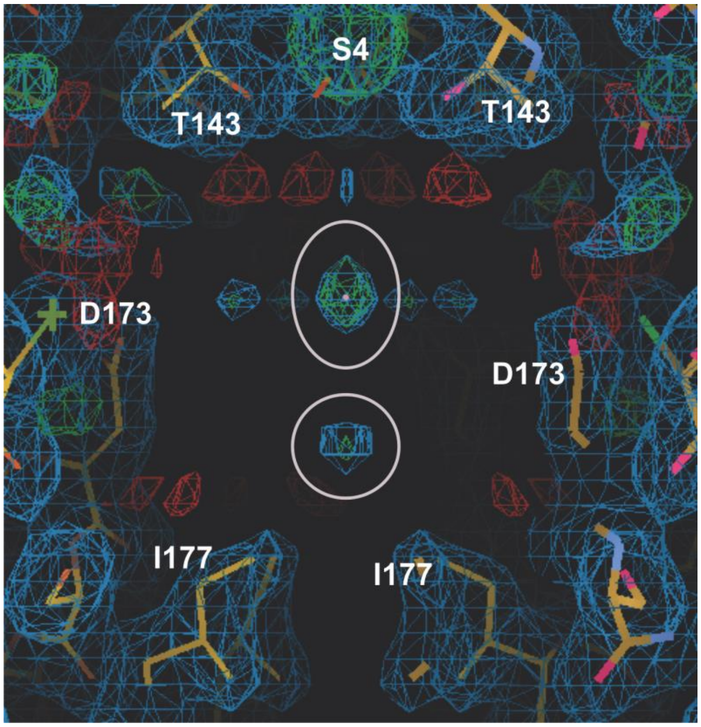


**Supplementary Figure S5. Electron density in the inner cavity of Apo-KW (5KUK) crystal diffracting to 2.0 Å.** 2fo-fc map in cyan is contoured at 1 σ while fo-fc maps (green or red) are contoured at 3 σ. Gray circles indicate putative $K^+$ ion densities.

### Supplementary Movies

#### Movie 1. $K^+$ ion permeation through an open PIP2-KW/GD channel

KW/GD protein is shown in cartoon with the nearest subunit omitted for clarity. The residues forming the HBC are explicitly shown in sticks (I177, M181). Water molecules are shown in sticks, and $K^+$ ions are represented as gray spheres with one $K^+$ ion highlighted in yellow illustrating complete $K^+$ ion permeation event starting from the bottom entrance to the extracellular side of the protein.

#### Movie 2. $K^+$ ion permeation through an open PIP2-KW/GD(G) channel

A similar movie illustrating complete $K^+$ ion conduction through KW/GD(G) structure.

**Movie 3. Gating transition from a closed to an open state**. $PIP_2$-KW crystal structure is taken as closed state while the last snapshot of 1 µs $PIP_2$- KW/GD(G) MD simulation is taken as open state. The two structures are aligned to maximize overlap of the SF backbone atoms (142-147). The closed state structure is shown silver, and the structures in transition from the closed to the open state are shown in light green.

#### Movie 4. $K^+$ ion permeation at the selectivity filter

For better visibility, only two opposing subunits of the Kir2.2 are represented as sticks. $K^+$ ions are represented as spheres. Ions that are already within, or which enter, the selectivity filter are colored consecutively, from silver via bronze, gold and brown to black. Ions that are close to the selectivity filter but do not enter it within the simulation are colored white. Water is represented as sticks. Although ion binding site S4 is occasionally occupied by water, only $K^+$ ions in this site lead to subsequent ion advancement through the filter.
